# *Salmonella enterica* serovar Typhimurium ST313 sublineage 2.2 has emerged in Malawi with a characteristic gene expression signature and a fitness advantage

**DOI:** 10.1101/2023.07.11.548493

**Authors:** Benjamin Kumwenda, Rocío Canals, Alexander V. Predeus, Xiaojun Zhu, Carsten Kröger, Caisey Pulford, Nicolas Wenner, Lizeth Lacharme Lora, Yan Li, Siân V. Owen, Dean Everett, Karsten Hokamp, Robert S. Heyderman, Philip M. Ashton, Melita A. Gordon, Chisomo L. Msefula, Jay C. D. Hinton

## Abstract

Invasive non-typhoidal *Salmonella* (iNTS) disease is a serious bloodstream infection that targets immune-compromised individuals, and causes significant mortality in sub-Saharan Africa. *Salmonella enterica* serovar Typhimurium ST313 causes the majority of iNTS in Malawi, and we performed an intensive comparative genomic analysis of 608 isolates obtained from fever surveillance at the Queen Elizabeth Hospital, Blantyre between 1996 and 2018. We discovered that following the upsurge of the well-characterised *S.* Typhimurium ST313 lineage 2 from 1999 onwards, two new multidrug-resistant sublineages designated 2.2 and 2.3, emerged in Malawi in 2006 and 2008, respectively. The majority of *S.* Typhimurium isolates from human bloodstream infections in Malawi now belong to sublineage 2.2 or 2.3. To identify factors that characterised the emergence of the prevalent ST313 sublineage 2.2, we performed genomic and functional analysis of two representative strains, D23580 (lineage 2) and D37712 (sublineage 2.2). Comparative genomic analysis showed that the chromosome of ST313 lineage 2 and sublineage 2.2 were broadly similar, only differing by 29 SNPs and small indels and a 3kb deletion in the Gifsy-2 prophage region that spanned the *sseI* pseudogene. Lineage 2 and sublineage 2.2 have unique plasmid profiles that were verified by long read sequencing. The transcriptome was initially explored in 15 infection-relevant conditions and within macrophages. Differential gene expression was subsequently investigated in depth in the four most important *in vitro* growth conditions. We identified up-regulation of SPI2 genes in non-inducing conditions, and down-regulation of flagellar genes in D37712, compared to D23580. Following phenotypic confirmation of transcriptional differences, we discovered that sublineage 2.2 had increased fitness compared with lineage 2 during mixed-growth in minimal media. We speculate that this competitive advantage is contributing to the continuing presence of sublineage 2.2 in Malawi.

## Introduction

Non-typhoidal *Salmonella* (NTS) is a major pathogen that threatens people across the world. Typhimurium and Enteritidis are the two serovars of *Salmonella enterica* that cause the highest levels of self-limiting gastrointestinal disease in Europe, the USA and other high-income countries (Zhang *et al*., 2003). In the industrialised world, NTS has largely been associated with intensive food production, animal husbandry, and global distribution systems (Majowicz *et al*., 2010). Globally, the most common sequence type of *S*. Typhimurium associated with gastroenteritis is ST19. Diarrhoeal NTS disease (dNTS) is mainly foodborne and poses a significant burden to public health globally, causing approximately 153 million cases and 57,000 deaths per annum (Kirk *et al*., 2015; Chirwa *et al*., 2023).

In contrast, a lethal systemic disease called invasive non-typhoidal Salmonellosis (iNTS) has emerged in recent decades in low- and middle-income countries in sub-Saharan Africa. Cases of iNTS are characterized by bloodstream infections of immune-compromised individuals such as children under five years of age, and HIV-positive adults. Anaemia, malnutrition and malaria are some of the major risk factors (Feasey *et al*., 2012). In some countries of sub-Saharan Africa, *Salmonella* causes more cases of community-onset bloodstream infections than any other bacterial pathogen (Marchello *et al*., 2019). In 2017, 535,000 cases of iNTS disease were estimated worldwide, with about 80% of cases and 77,000 deaths occurring in sub-Saharan Africa (Stanaway *et al*., 2019)

Clinically, the treatment of iNTS is complicated by multi-drug (MDR) resistance which limits therapeutic options (Crump *et al*., 2015). Widespread resistance of iNTS pathogens to first-line drugs such as chloramphenicol, ampicillin and cotrimoxazole has been seen in many countries (Kariuki *et al*., 2006). This MDR phenotype may be one of the reasons the case fatality rate associated with iNTS is amongst the highest in comparison to any infectious disease (15%) (Marchello *et al.,* 2022) Resistance to second line drugs such as ceftriaxone, ciprofloxacin and azithromycin has been reported in a few African countries (Tack *et al*., 2020). Clearly, the problem of MDR *Salmonella* must be addressed urgently (Gilchrist and MacLennan, 2019).

The African iNTS epidemic is mainly caused by two *Salmonella* pathovariants, *S.* Typhimurium sequence type 313 (ST313) and specific clades of *S.* Enteritidis (Kingsley *et al*., 2009; Okoro *et al*., 2012; Feasey *et al*., 2016). *S.* Typhimurium ST313 is responsible for about two-thirds of clinical iNTS cases that have been reported in Africa (Gilchrist and MacLennan, 2019).

It is not certain how these pathogens are transmitted, but there is increasing evidence from case-control studies that ST313 strains are human-associated but not animal-associated within households (Post *et al*., 2019; Koolman *et al*., 2022). A recent summary concludes that the available data are consistent with the person-to-person transmission hypothesis for iNTS disease (Chirwa *et al*., 2023). Global efforts to combat iNTS infections are currently focused on vaccine development which is currently progressing to clinical trials (Piccini and Montomoli, 2020).

Since 1998, continuous sentinel surveillance for fever and bloodstream infections among adults and children has been undertaken at Queen Elizabeth Central Hospital (QECH). This tertiary referral hospital in Blantyre, Malawi, serves an urban population of about 920,000 with a high incidence of malaria, HIV and malnutrition (Musicha *et al*., 2017). Following blood-culture of samples collected from patients of all ages presenting with fever, whole genome sequencing identified the ST313 variant of *S.* Typhimurium (Kingsley *et al*., 2009). Phylogenetic analysis revealed that the chloramphenicol-sensitive ST313 lineage 1was clonally-replaced in Malawi by the chloramphenicol-resistant lineage 2 (Okoro *et al*., 2012). More recently, a ST313 sublineage II.1 (2.1) emerged from lineage 2 in Democratic Republic of Congo (DRC) in Central Africa. Sublineage 2.1 had altered phenotypic properties including biofilm formation and metabolic capacity and resistance to azithromycin (Van Puyvelde *et al*., 2019).

An initial suggestion that ST313 lineage 2 was undergoing evolutionary change in East Africa came from a small study that identified seven *S.* Typhimurium ST313 Malawian isolates, dated between 2006 and 2008, that differed from lineage 2 by 22 core-genome single nucleotide polymorphisms (SNPs) (Msefula *et al*., 2012).

To begin to examine the evolutionary trajectory of *S.* Typhimurium in Malawi at a large scale, we conducted a comparative genomic analysis study focused on 680 isolates dating between 1998 and 2018 (Pulford *et al*., 2021). We previously confirmed that ST313 lineage 1 (L1) was replaced by lineage 2 (here designated L2.0), and discovered an antibiotic-sensitive lineage 3 (L3) that emerged in 2016 (Pulford *et al*., 2021).

We performed a more intensive phylogenetic analysis of the same collection of *S.* Typhimurium ST313 isolates, most of which caused bloodstream infections in Malawi over two decades. We discovered two novel sublineages named 2.2 (L2.2) and 2.3 (L2.3) that have been replacing L2.0 since 2006.

Here we present a comprehensive comparative genomic analysis of the most prevalent ST313 L2.2 sublineage, and report the results of a functional genomic approach that identified key phenotypic characteristics that distinguish L2.2 from L2.0.

## Results

### Identification of *S.* Typhimurium ST313 sublineages 2.2 and 2.3 in Malawi

The *S.* Typhimurium ST313 L2 (Lineage II) was originally identified as the major cause of iNTS cases across sub-Saharan Africa in the early 2000’s (Kingsley *et al*., 2009; Okoro *et al*., 2012) (Okoro *et al*., 2015). Subsequently, an azithromycin-resistant variant of *S.* Typhimurium ST313 was found in a single country, the Democratic Republic of Congo between 2008 and 2016, and was designated sublineage L2.1 (Van Puyvelde *et al*., 2019).

To investigate the evolutionary dynamics of *S.* Typhimurium ST313 L2 in Malawi over a 22 year period, we focused on the large collection of 8,000 *S.* Typhimurium isolates derived from bloodstream infection in hospitalised patients at the Queen Elizabeth Central Hospital, Blantyre, Malawi (Feasey *et al.,* 2015). The collection was assembled by the Malawi–Liverpool– Wellcome Trust Clinical Research Programme (MLW) between 1996 and 2018; the precise annual numbers of isolates are shown in Fig 1B. A random sub-sampling strategy was used to select 608 isolates selected for whole-genome sequencing which included 549 *S.* Typhimurium ST313 isolates (Pulford *et al*., 2021).

**Fig 1.**
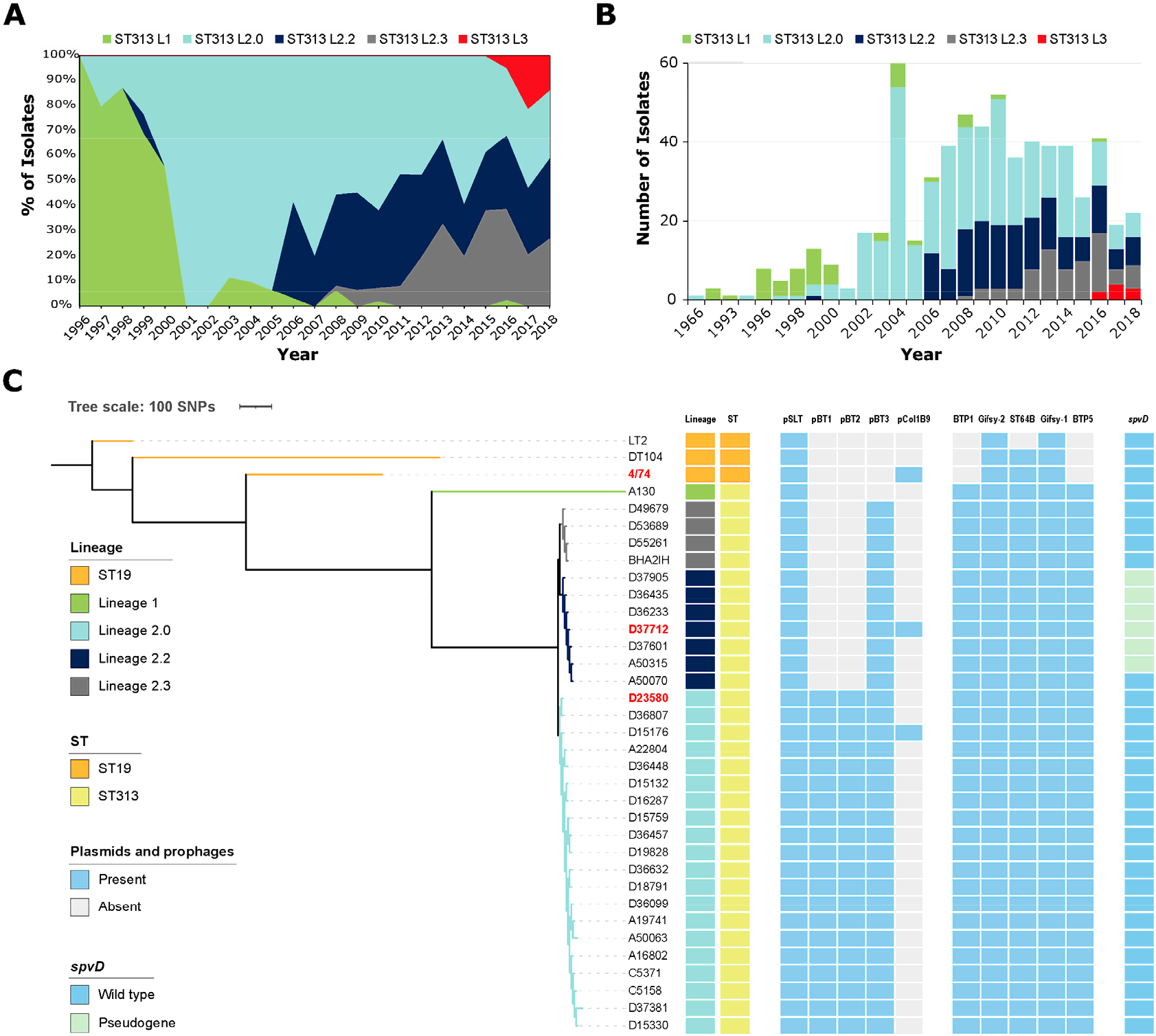
Emergence of *S.* Typhimurium ST313 sublineages L2.2 and L2.3 in Malawi. **(A)** Evolutionary dynamics of S. Typhimurium lineages in Blantyre, Malawi from 1996 to 2018. The genomes of 549 *S*. Typhimurium ST313 isolates from bacteraemic patients at the Queen Elizabeth Hospital in Blantyre, Malawi were used for this analysis. The proportions of the five lineages/sublineages are shown. **(B)** The total number of isolates of each lineage/sublineage per year. **(C)** Phylogenetic comparison between representative strains of *S.* Typhimurium ST19 and four ST313 lineages/sublineages (L1, L2.0, L2.2, L2.3) showing the presence and absence of plasmids, prophages and the *spvD* pseudogene. The complete phylogenetic analysis of 707 *S.* Typhimurium genomes is shown in Fig. S1.

Here, we used a core-gene SNP-based maximum likelihood (ML) phylogenetic tree to investigate the population structure of *S.* Typhimurium ST313 L2.0 in more detail (Fig. S1). As well as identifying members of the antibiotic-sensitive lineage 3 that we reported previously (Pulford *et al*., 2021), we discovered that ST313 L2 could be split into three phylogenetically-distinct sublineages that differed by 39 SNPs. The *S.* Typhimurium ST313 L2 reference strain D23580 (Kingsley *et al*., 2009) belonged to the first sublineage, which we have now designated as ST313 L2.0 (Fig 1C). As ST313 sublineage L2.1 has been defined previously (Van Puyvelde *et al*., 2019), the new sublineages were designated as L2.2 and L2.3, and belonged to different hierBAPS level 2 clusters (Fig 1C and Fig S1). We identified 151 L2.2 isolates and 74 L2.3 isolates, against a backdrop of 350 L2.0 isolates.

In Blantyre, Malawi, *S.* Typhimurium ST313 L2.2 was first detected in 2006, and L2.3 was initially observed in 2008 (Fig. 1A. Both L2.2 and L2.3 increased in prevalence at the Queen Elizabeth Central Hospital in Blantyre in subsequent years. By 2018, L2.2 and L2.3 had largely replaced L2.0 (Fig 1A-B). Our published Bayesian (BEAST) analysis (Pulford *et al*., 2021) estimated that the Most Recent Common Ancestor (MRCA) of ST313 lineage 2 dates back to 1948 (95% HPD = 1929-1959).

To understand the accessory gene complement of L2.2 and L2.3, we compared the genomes of seven L2.2 isolates and four L2.3 isolates with 17 L2.0 isolates, ST313 L1 and ST19 (Fig 1C, Table S1). *S.* Typhimurium strain D23580 is the representative strain of L2.0 (Kingsley *et al*., 2009), for which we previously used long-read sequencing and other approaches to thoroughly characterise the chromosome and the plasmid complement (Canals *et al*., 2019b).

### Antimicrobial Resistance

AMR variants of *S.* Typhimurium with resistance to ampicillin and cotrimoxazole were detected at an early stage of the iNTS epidemic, from 1997 onwards (Gordon *et al.,* 2008). Multidrug-resistant variants of *S.* Typhimurium ST313 that were no longer susceptible to chloramphenicol, ampicillin and cotrimoxazole subsequently emerged in Malawi (Gordon *et al.,* 2008) and have been reported elsewhere in sub-Saharan Africa by the GEMS study (Kasumba *et al*., 2021). The *S.* Typhimurium ST313 L2.0, L2.2 and L2.3 isolates shared the same MDR profiles (resistance to chloramphenicol, ampicillin and cotrimoxazole), and carried identical IS21-AMR gene cassettes within the pSLT-BT plasmid.

### Comparative genomics of *S.* Typhimurium ST313 sublineage 2.2

Because *S.* Typhimurium ST313 L2.2 was the predominant novel sublineage in Blantyre, Malawi, we focused on L2.2 for the remainder of this study. We used the phylogeny (Fig 1C) to select strain D37712 as a representative isolate of L2.2. D37712 was isolated from the blood of an HIV-positive Malawian male child and has been deposited in the National Collection of Type Cultures (NCTC). The initial genome sequence of D37712 was obtained in 2012 with Illumina technology, an assembly that comprised 27 individual contigs (Msefula *et al*., 2012). To generate a reference-quality genome, we resequenced D37712 with both long-read PacBio and Illumina short-read technologies. Our hybrid strategy generated a complete genome assembly that included one circular chromosome and three plasmids (see Materials & Methods; GenBank CP060165, CP060166, CP060167 and CP060168). This high-quality genome sequence allowed us to conduct a detailed comparative genomic analysis of L2.2 strain D37712 with L2.0 strain D23580 (accession number FN424405), summarised in Fig. 2 and Table S2.

**Fig 2.**
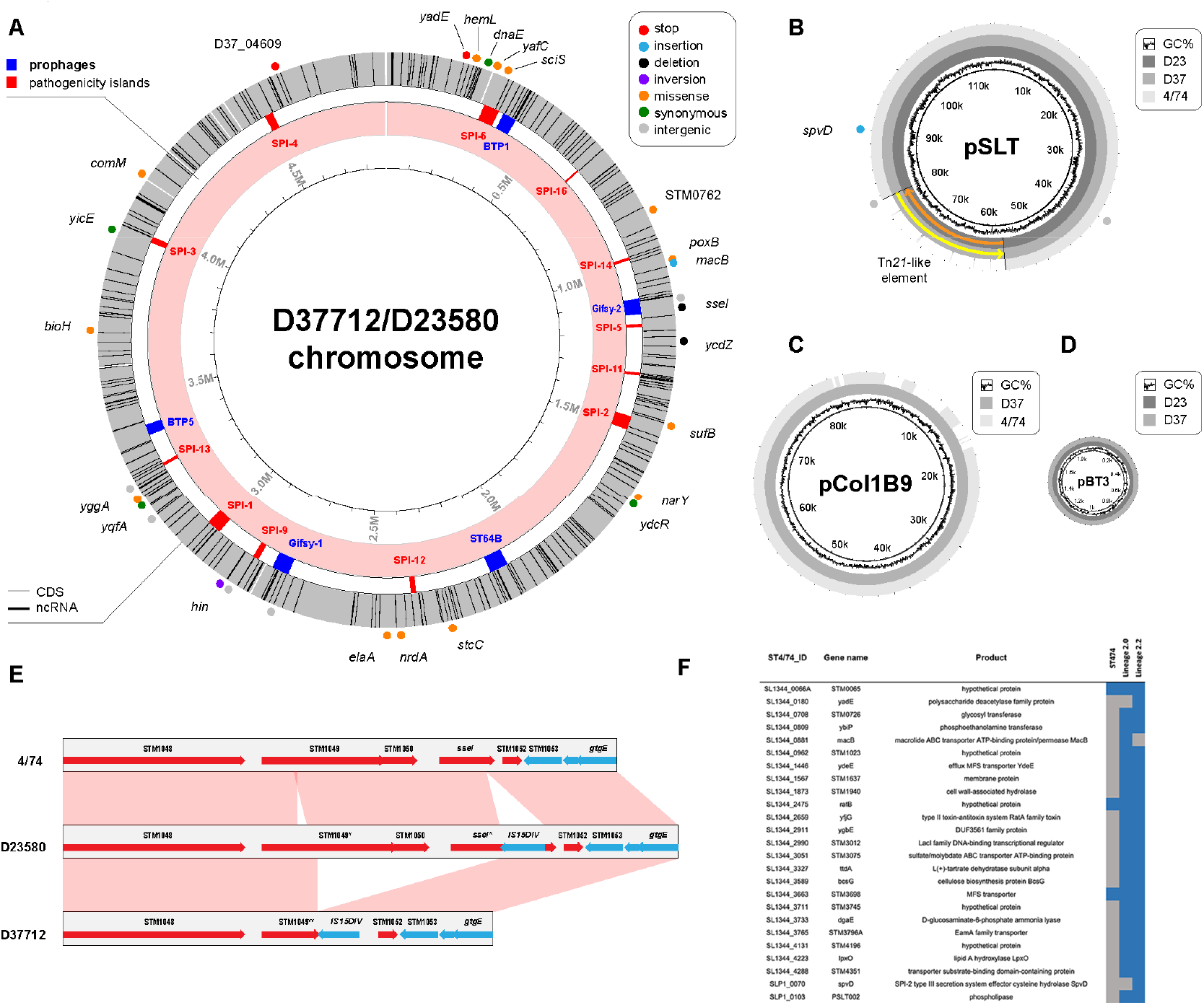
Key genetic similarities and differences between the chromosome and plasmid profiles of D23580 (lineage 2) and D37712 (L2.2). **(A)** A comparison of the D23580 (L2.0) and D37712 (L2.2) chromosomes. The dots around the chromosome are different kinds of SNPs identified. Phages and *Salmonella* pathogenicity islands are shown in blue and red respectively. **(B)** Plasmid profile of D37712 versus D23580. The pSLT-BT virulence plasmid is present in both D37712 and D23580, and carries the Tn-21 transposable element; **(C)** pCol1B9 is present in D37712 and absent from D23580 (D) pBT3 is present in both D37712 and D23580. **(E)** Absence of *sseI* gene and the STM1050 coding sequence in L2.2 (D37712), as compared to *S.* Typhimurium ST19 4/74 and *S.* Typhimurium ST313 L2.0 (D23580). **(F)** List of pseudogenes in D37712 and D23580, with reference to 4/74. The colour blue means pseudogene/disrupted gene while grey indicates functional genes. *macB* is a pseudogene in D23580 (L2.0) but not in L2.2, while *spvD* is a pseudogene in L2.2 but not in L2.0. All L2.2 strains share similar pseudogenes.

Overall, the two strains contain a similar number of genes. The D37712 and D23580 genomes shared 5,016 orthologous genes, including 4,729 protein-coding genes and pseudogenes as well as the 287 small RNA (sRNA) genes that we identified previously. The D23580 annotation contains 4,823 protein-coding and pseudogenes and 287 sRNAs (Canals *et al.,* 2019b), while D37712 contains 4,821 protein-coding and pseudogenes and 287 sRNAs.

### Overview of D23580 and D37712 genomes

The chromosomes of D23580 and D37712 are 4,879,402 and 4,876,060 bp, respectively, and similar in size to other S. Typhimurium genomes (Kingsley *et al*., 2009; Branchu *et al*., 2018). The D23580 and D37712 strains share a similar prophage profile, with both strains carrying five prophages (BTP1, Gifsy-2, ST64B, Gifsy-1, and BTP5) which were located at the same positions on the chromosome. Previously, we have established that just one of these prophages, BTP1, is functional (Owen *et al*., 2017). The BTP1 prophage of D23580 encodes the novel BstA phage defence system (Owen *et al.,* 2021) and a particularly high level of viable BTP1 phages is produced by spontaneous induction (Owen *et al*., 2017).

### Comparison of D23580 and D37712 chromosomes

The detailed genomic comparison of D37712 with D23580 showed that the two genomes were remarkably similar. Overall, the only differences between the genomes of the L2.0 and L2.2 strains were 26 chromosomal SNPs and small indels, plus one large deletion, and an inversion of the *hin* switch. In-depth annotation of the nucleotide variants identified 3 putative loss-of-function mutations (2 stop mutations, 1 frameshift insertion), 1 disruptive in-frame deletion, 4 synonymous mutations, 13 missense mutations, and 5 intergenic variants, summarised in Fig 2A.

The 3,358 bp-long deletion of a Gifsy-2 prophage-associated region that spanned the *sseI* pseudogene of D23580 removed two coding sequences (STM1050-51; STMMW_10611-STMMW_10631), and substantially truncated the STM1049 (STMMW_10601) gene (Fig 2E). The *sseI* gene encodes a cysteine hydrolase effector protein that modulates the directional migration of dendritic cells during systemic infection (Brink *et al*., 2018). In strain D23580, the insertion of a transposable element IS15DEV inactivated the *sseI* gene (Kingsley *et al*., 2009) causing increased dendritic cell-mediated dissemination of strain D23580 during infection (Carden *et al*., 2017). To confirm that the 3,358 bp deletion removed the *sseI* gene from the chromosome of strain D37712, we used an independent PCR-based approach (Fig S2).

### Comparison of D23580 and D37712 plasmids

ST313 L2.0 strain D23580 carries four plasmids, pSLT-BT, pBT1, pBT2 and pBT3 (Kingsley *et al*., 2009). In contrast, ST313 L2.2 carried a distinct plasmid complement (Fig 1C, Fig. 2BCD). In summary, strain D37712 carried pSLT-BT, pBT2 and pCol1B9 as detailed below. Both strains had a variant of the pSLT-BT virulence plasmid (Kingsley *et al*., 2009) that contains a Tn21-like transposable element with five antibiotic resistance genes. The D37712 version of pSLT-BT is similar to that of D23580, with two important differences (Fig 2B). Firstly, the Tn21-like element is inserted in the opposite direction with regards to the rest of the plasmid, suggesting that the transposable element remains active. Secondly, three nucleotide variants were identified in the pSLT-BT variant, two deletions in noncoding regions, and one frameshift insertion that generates a pseudogene of *spvD*. The SpvD effector protein, a cysteine protease, is translocated by the SPI2 type 3 secretion system and suppresses the NF-κB-mediated pro-inflammatory immune response and contributes to virulence in mice (Grabe *et al*., 2016).

Plasmid pCol1B9 was of particular interest because it was absent from D23580, but is present in *S.* Typhimurium ST19 strain 4/74 (Richardson *et al*., 2011). 4/74 is the parent of the *S.* Typhimurium SL1344 strain that has been used extensively for the study of *S*. Typhimurium pathogenesis and gene regulation since 1986 (Kröger *et al*., 2012; Rankin & Taylor, 1966). Our annotation of the pCol1B9 plasmid included 95 distinct protein-coding genes, while the previously published annotation of pCol1B9^4/74^ assigned 101 protein-coding genes. Some of these represent annotation discrepancies, while others represent true genetic differences (Fig. S3). Upon careful examination, 14 genes were unique to pCol1B9^D37712^, while 20 were unique to pCol1B9^4/74^. There were 81 genes carried by both plasmids. Interestingly, pCol1B9^D37712^ lacked the colicin toxin-antitoxin system that both gave pCol1B9 its name, and provides *Salmonella* with a competitive advantage in the gut (Nedialkova *et al*., 2014). The pCol1B9^D37712^ plasmid carried a locus that was absent from pCol1B9^4/74^, namely the *impC-umuCD* operon (Fig. S3) which encodes the error-prone DNA polymerase V responsible for the increased mutation rate linked to the SOS stress response in *E. coli* (Sikand *et al*., 2021).

An 85 kb plasmid carried by D23580, pBT1, was previously shown by our laboratory to play an important role in *Salmonella* biology by encoding an orthologous *cysS* gene responsible for expressing the essential cysteinyl tRNA-synthetase enzyme (Canals *et al.,* 2019b). This pBT1 plasmid was completely absent from D37712, and from all isolates of sublineage L2.2 that were examined (Fig. 1C).

### Comparison of pseudogene status of D23580 and D37712

Our comparative genomic analysis focused on the pseudogenes found in strains 4/74, D23580, and D37712 (Fig 2F, Table S3). The pseudogenisation of several D23580 genes, compared with strain 4/74, have been linked to the invasive phenotype of African *Salmonella* ST313 (Kingsley *et al*., 2009). We found that the pseudogene complement of D23580 was largely conserved in D37712. We have recently reported the role of the MacAB-TolC macrolide efflux pump in the virulence of *S.* Typhimurium ST313, and showed experimentally that *macB* was an inactive pseudogene in D23580 (Honeycutt *et al*., 2020). Interestingly, the *macB* gene is functional in D37712. Compared with D23580, three additional D37712 genes were pseudogenised (*spvD, yadE,* and STMMW_42692).

Overall the chromosomes of ST313 lineage 2 and sublineage 2.2 were highly-conserved and differed by just 29 SNPs/ small indels, and a 3kb deletion in the Gifsy-2 prophage region. The ST313 lineage 2 and sublineage 2.2 have distinct plasmid profiles.

### Transcriptional landscape of S. Typhimurium ST313 sublineage L2.2

Previously, we characterized the primary transcriptome of two other *S.* Typhimurium strains, 4/74 and D23580, using a combination of multi-condition RNA-seq and differential RNA-seq (dRNA-seq) techniques (Canals *et al*., 2019b; Kröger *et al*., 2013). To identify the transcriptional start sites (TSS) of strain D37712, we analysed a pooled sample containing RNA from 15 *in vitro* conditions by dRNA-seq and RNA-seq as detailed previously (Kröger *et al*., 2013). The high similarity between the D23580 and D37712 chromosomes allowed us to map the curated set of TSS that were previously defined for D23580 (Hammarlof *et al.,* 2018) onto a combined D37712/D23580 reference genome. To allow individual TSS to be examined in particular chromosomal or plasmid regions, data from both the dRNA-seq and pooled RNA-seq experiments can be visualised in our online genome browser (http://hintonlab.com/jbrowse/index.html?data=Combo_D37/data).

### Preliminary gene expression profiling of S. Typhimurium ST313 sublineage L2.2

Given the high level of similarity between the genomes of L2.2 and L2.0, we went on to identify differences at the transcriptional level. We performed a multi-condition RNA-seq-based transcriptomic analysis of gene expression profiles of L2.2 strain D37712 without biological replicates.

This comparative transcriptomic screen was based on our published approach (Canals *et al*., 2019b). Specifically, we used 15 individual infection-relevant *in vitro* conditions (Kröger *et al*., 2013) and did intra-macrophage transcriptome profiling using the protocol previously established for *S.* Typhimurium ST19 (Srikumar *et al*., 2015). The RNA-seq samples were mapped to a combined reference genome, which included the annotated D23580 chromosome (Canals *et al*., 2019b), as well as all the plasmids described earlier (pSLT-BT, pBT1, pBT3 and pCol1B9; see Methods). The initial RNA-seq assessment (detailed in Methods) involved 2-4M non-rRNA/tRNA reads per sample, allowing gene signatures specific for each *in vitro* condition to be identified. Although single replicate RNA-seq experiments of this type cannot be used for statistically-robust differential gene expression analysis, they do provide a useful screening approach for identifying growth conditions to be used for follow-up experiments. The individual RNA-seq experiments showed broad condition-specific similarities in gene expression between strains 4/74, D37712, and D23580 (Fig 3A). The gene expression values from each profiled condition are available as raw counts and TPMs in Tables S4 and S5.

**Fig 3.**
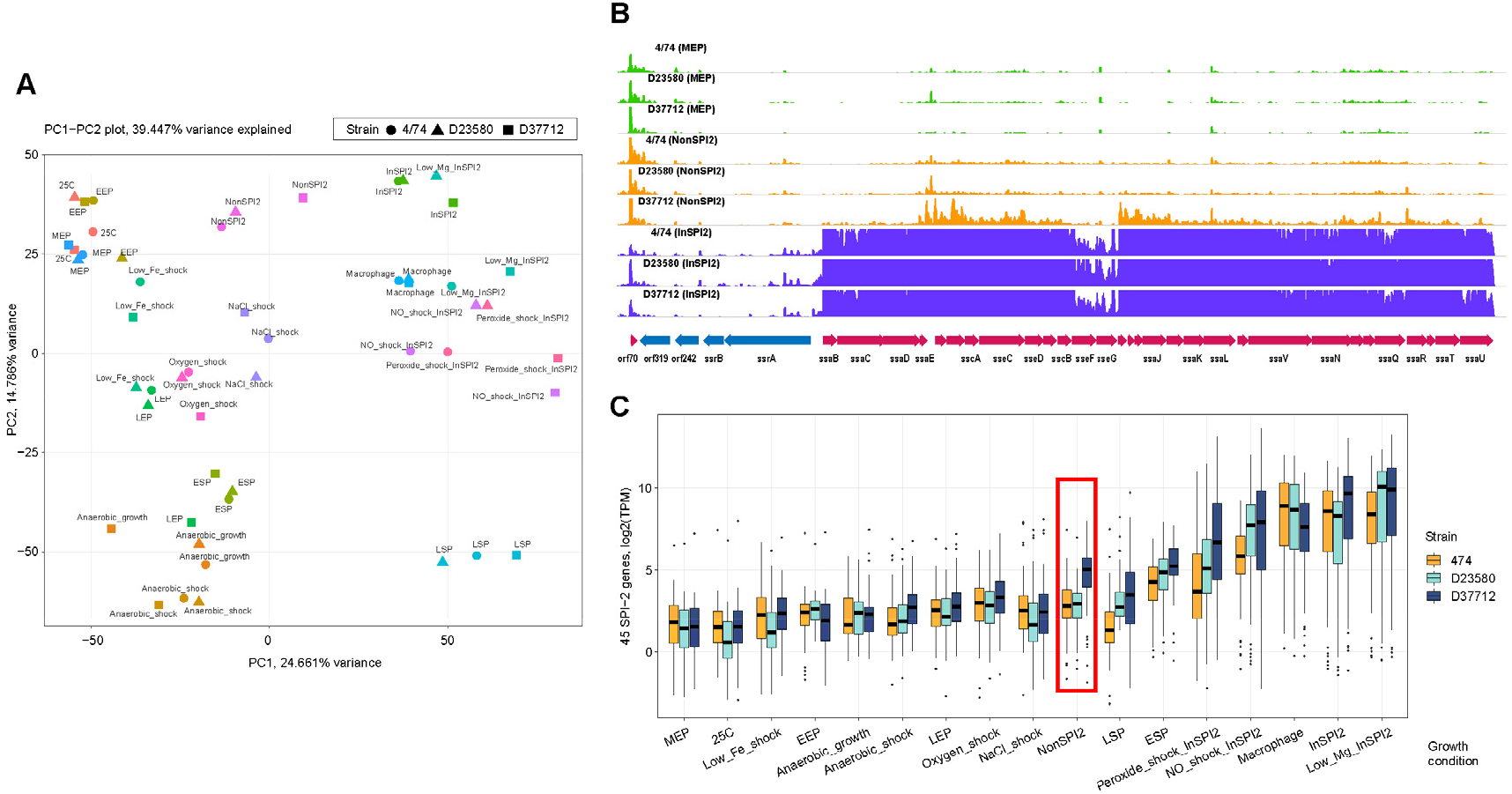
General comparison of expression profiles of strains 4/74, D23580, and D37712 under 17 different *in vitro* conditions. **(A)** Principal component analysis (PCA) plot of the individual RNA-seq samples, indicating the overall similarity in gene expression between the three strains. The 17 growth conditions have been defined previously (Kröger *et al*., 2013). **(B)** Visualization of SPI-2 pathogenicity island expression with the Jbrowse genomic browser, under mid-exponential phase (MEP), InSPI2, and NonSPI2 *in vitro* conditions. **(C)** Boxplot visualization of SPI-2 gene expression under mid-exponential phase (MEP), InSPI2, and NonSPI2 *in vitro* conditions. The elevated expression of SPI-2 genes in strain D37712 cultured under NonSPI2 conditions is highlighted in a red box.

To select the ideal environmental conditions to use for subsequent experiments, we assessed the expression profiles of known *Salmonella* pathogenicity islands which were broadly similar in strains D37712, and D23580. Although the expression profile of the SPI2 pathogenicity island was broadly similar between D37712, D23580 and 4/74 in most growth conditions, the SPI2 genes of D37712 were highly up-regulated in a single growth condition, NonSPI2 (Fig. 3B-C). NonSPI2 is a minimal medium with a neutral pH and a relatively high level of phosphate, in which S. Typhimurium does not usually express the SPI2 pathogenicity island (Löber *et al*., 2006; Kröger *et al*., 2013). This intriguing observation prompted us to perform the more discriminating set of transcriptomic experiments described below.

### Differential gene expression analysis of *S*. Typhimurium D37712 versus D23580 in four *in vitro* conditions with multiple biological replicates

To define the transcriptional signature of strain D37712 more accurately, we generated RNA-seq data from D37712 grown in four *in vitro* conditions that stimulate expression of the majority of virulence genes: ESP, anaerobic growth, NonSPI2 and InSPI2, with multiple (3-4) biological replicates. We compared the results with our published transcriptomic data for S. Typhimurium strains 4/74 and D23580 (Canals *et al.,* 2019b; Kröger *et al*., 2013). Differential expression analysis with DEseq2, with conservative cut-offs (fold change ≥ 2, FDR ≤0.001), showed that the gene expression profiles of D37712 and D23580 were broadly similar, and shared key differences to the transcriptional profile of strain 4/74 under each of the four *in vitro* conditions (Fig 4A). The differential expression results are summarized in Table S6.

**Fig 4.**
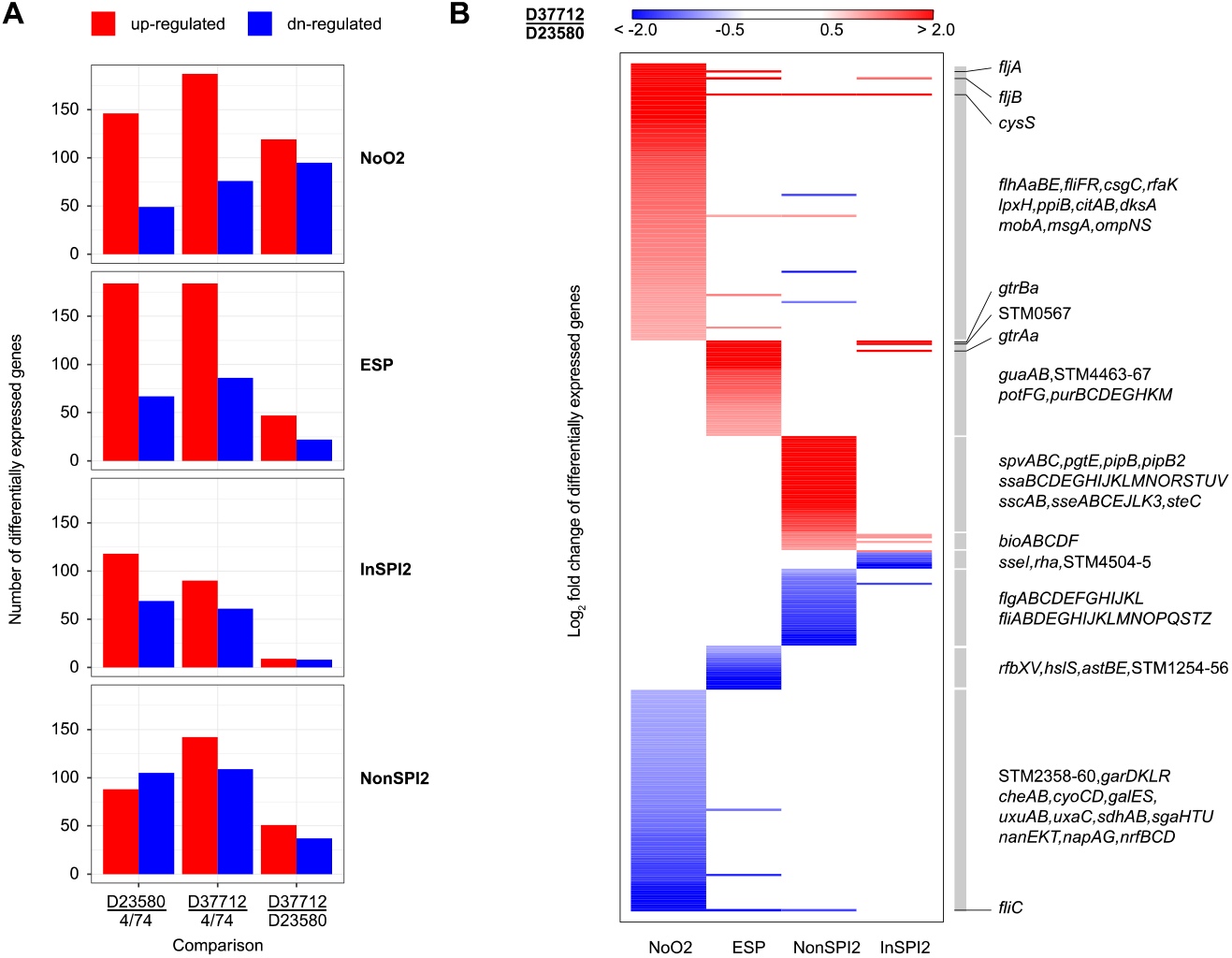
Differential gene expression of *S.* Typhimurium 4/74, D37712, and D23580 under 4 in vitro conditions. (**A)** Boxplots indicating the number of differentially-expressed genes identified in the following *in vitro* growth conditions: early stationary phase, ESP; anaerobic growth, NoO2; SPI-2 inducing medium, InSPI2; SPI-2 non-inducing minimal medium, NonSPI2. Multiple (3 to 5) biological replicates were used for comparison. DESeq2 was used for differential analysis; only genes with |log2FC| ≥ 1 and with adjusted *p*-value ≤ 0.001 were retained. **(B)** Heatmap of the genes differentially expressed between D23580 and D37712. Functional groups and operons of interest are highlighted on the right of Panel B.

We specifically investigated transcription of the *pgtE* gene, which encodes the outer-membrane protease previously linked to the ability of African *Salmonella* ST313 to resist human serum killing (Hammarlöf *et al*., 2018). Compared to 4/74, the *pgtE* gene of both the D23580 and D37712 strains showed a similar pattern of up-regulation by a factor of 7 to 18 across all conditions. This finding is consistent with the fact that D37712 carries the same T nucleotide in the -10 region of the *pgtE* promoter that is responsible for increased expression of the *pgtE* transcript in strain D23580 (Hammarlöf *et al*., 2018).

The majority (92%) of 4,729 orthologous coding genes of both D37712 and D23580 were expressed at similar levels. We identified a total of 364 genes that were differentially expressed in at least one growth condition between D37712 and D23580 as follows: ESP (69 differentially-expressed genes), anaerobic growth (214 differentially-expressed genes), NonSPI2 (88 differentially-expressed genes) and InSPI2 (17 differentially-expressed genes; Fig 4B).

Overall, the differentially expressed genes that distinguished D37712 from D23580 were seen in a single growth condition and included flagellar genes (down-regulated), SPI2-associated genes(up-regulated), and genes involved in general and anaerobic metabolism (down-regulated).

The SPI2 pathogenicity island genes play a key role in the intracellular replication of *S.* Typhimurium, and encode the type III secretion system that is responsible for translocation of key effector proteins into mammalian cells (Jennings *et al*., 2017). The RNA-seq data showed that SPI2 genes were expressed at similarly high levels in both D37712 and D23580 strains following induction (InSPI2 media; Fig 4B), and confirmed that the key SPI2 expression difference was only seen in strain D37712 under non-inducing growth conditions (NonSPI2 media). It is important to put this differential SPI2 expression into context. D37712 expresses SPI2 genes at about a 10-fold higher level than D23580 during growth in non-inducing NonSPI2 media, but the actual level of expression was 20-fold less than the level stimulated by growth in SPI2-inducing conditions (InSPI2 medium).

The up-regulation of *fljA* and *fljB* and the down-regulation of *fliC* in D37712, compared to D23580 in all four growth conditions likely reflects the opposite orientation of the *hin* switch in the D37712 genome compared to D23580. This type of *hin* inversion occurs frequently in *S.* Typhimurium (Johnson and Simon, 1985).

Another gene that was up-regulated in D37712 across all profiled conditions was the chromosomally-encoded *cysS^chr^*, that encodes cysteine-tRNA synthetase. Previously, we reported that transcription of the *cysS^chr^* of strain D23580 was uniformly down-regulated compared to 4/74, a defect that was compensated by the presence of a pBT1 plasmid-encoded cysteine-tRNA synthetase (Canals *et al.,* 2019a). Increased expression of the chromosomal *cysS* gene in D37712 was consistent with the absence of the pBT1 plasmid. Our comparative transcripomic analysis showed that expression levels of *cysS* were similar in D37712 and 4/74 under all growth conditions.

Numerous virulence genes and operons were differentially expressed between D23580 and D37712. The SPI-16-associated *gtrABCa* operon (STM0557, STM0558, STM0559) is responsible for adding glucose residues to the O-antigen subunits of LPS that enhance the long-term colonisation of the mammalian gastrointestinal tract by *S.* Typhimurium ST19 (Bogomolnaya *et al*., 2008). We found that the *gtrABCa* genes were significantly up-regulated in several conditions in D37712, compared to both D23580 and 4/74.

The *spvABCD* operon of D37712 was up-regulated under non-SPI2-inducing growth conditions, compared to D23580. A signature pseudogene of ST313 L2.2 is the frameshift insertion in the *spvD* gene that generates a truncated version of the SpvD protein. The H199I mutation at position 199 and the associated 17 amino acid truncation is predicted to ablate the activity of the SpvD cysteine protease (Grabe *et al*., 2016). The functional consequences of the *spvD* variant of ST313 L2.2 strain D37712 and the up-regulation of the *spvABCD* operon remain to be established experimentally.

### The SalComD37712 community transcriptional data resource

To allow scientists to gain their own biological insights from analysis of this rich transcriptomic dataset, the transcriptomic and gene expression data generated in this study are presented online in a new community resource, <u>SalComD37712</u>. The data resource shows the expression levels of all D37712 coding and non-coding genes, including both chromosomal and plasmid-encoded transcripts. The SalComD37712 website complements our existing SalComD23580 (https://tinyurl.com/SalComD23580) resource, and adds an inter-strain comparison of gene expression profiles between D37712 and D23580 as well as normalized gene expression values (TPM), using an intuitive heat map-based approach. <u>SalComD37712</u> included our published RNA-seq data (Canals *et al*., 2019b), re-analysed with an updated bioinformatic pipeline and a combined reference genome (see Methods). This online resource facilitates the intuitive interrogation of transcriptomic data as described previously (Perez-Sepulveda and Hinton, 2018).

Additionally, we generated a unified genome-level browser that provides access to the *S.* Typhimurium L2.2 D37712 transcriptome, in the context of our previously published RNA-seq data for the L2.0 strain D23580 and the ST19 strain 4/74. This novel “combo” browser is available at http://hintonlab.com/jbrowse/index.html?data=Combo_D37/data.

### Identification of phenotypes that distinguish ST313 sublineage L2.2 from L2.0

To explore the phenotypic impact of the transcriptomic signature of L2.2 (D37712), we performed a series of motility experiments, fluorescence-based gene expression experiments and mixed-growth assays.

D33712 showed a significantly decreased level of motility on NonSPI2 minimal media, compared with both the ST19 strain 4/74 and the L2 D23580 strain (Fig. 5A). This finding was consistent with the transcriptomic data, which showed down-regulation of D37712 flagellar genes compared with D23580 in the NonSPI2 condition (Fig. 4). In contrast, no differential expression of flagellar genes was seen between D33712 and D23580 in the InSPI2 growth condition (Fig. 4). The decreased motility phenotype may be linked to the inversion of the *hin* element detailed above. The flagella system encodes a distinct type III secretion apparatus responsible for the dual functions of bacterial motility and activation of the mammalian innate immune system via TLR5 (Lai *et al*., 2013).

**Fig 5.**
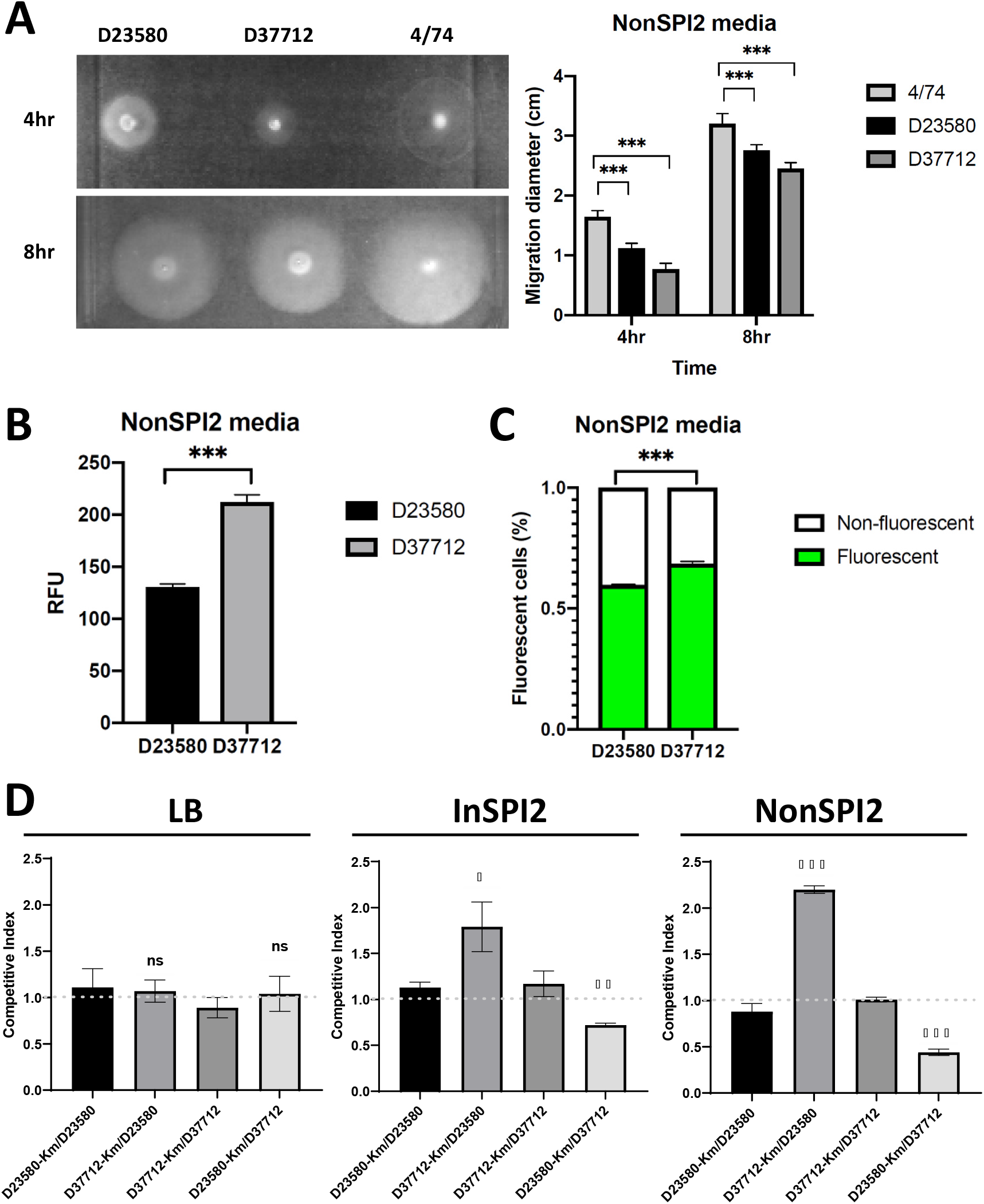
Phenotypes that distinguish ST313 L2.2 from ST313 L2.0. **(A)** Swimming motility assay of strains D23589, D37712 and 4/74, with a representative plate shown on the left. Average migration diameters were measured after 4 and 8 hours. Each bar represents the mean of three biological replicates, with *error bars* representing standard deviation. Significant difference (***) indicates *P* value (*t* test) < 0.001. In Panels B & C, comparison of *ssaG* expression by flow cytometry using D23580 and D37712 derivatives containing a chromosomal *ssaG*-GFP^+^ transcriptional fusion, strains SZS008 and SZS032, respectively. Cells were collected at 8 hours after inoculation in NonSPI2 media. Ten thousand events were acquired for each sample. **(B)** Mean fluorescent intensity signal of *ssaG*-GFP^+^ for D23580 (SZS008, dark grey) and D37712 (SZS032,, grey). Significant difference (***) indicates *P* value (*t* test) < 0.001. **(C)** Percentage of positive (green) and negative cells (white) for *ssaG* expression in each sample. Each bar represents the mean of three biological replicates, error bars show standard deviation. Significant difference (***) indicates *P* value (*t* test) < 0.001. **(D)** Relative fitness of wild-type D23580 and D37712 and their kanamycin resistant derivatives. Bacterial numbers were determined by overnight culture of a 1:1 mixture (wild-type versus Km^R^) in NonSPI2 (red), InSPI2 (blue) and LB (black) media. Each bar represents the mean of three biological replicates with *error bars* representing standard error. *P* values were determined by *t* test (***: *P* < 0.001; **: *P* < 0.01; *: *P* < 0.05; ns: no significance). A competitive index of 1 indicates the equal fitness of two strains, while a number higher than 1 reflects the increased fitness of kanamycin-resistant derivatives.

A key transcriptomic finding for strain D33712 was the expression of SPI2 genes during growth in an unusual environmental condition (NonSPI2) (Fig. 3B-C and Fig. 4B). NonSPI2 media differs from InSPI2 media by having a higher pH (pH7.4 versus pH5.8) and a higher level of phosphate (Löber *et al*., 2006). This apparent differential expression of SPI2 genes at the transcriptomic level under non-inducing conditions led us to investigate the expression of SPI2 at a single cell level using fluorescence transcriptional fusions. First, we introduced an *ssaG*-GFP^+^ transcriptional fusion into the chromosome of strains D33712 and D23580 (Methods; Table S8) to interrogate expression of the key SPI2 operon with flow cytometry. Figure 5B shows that in NonSPI2 media, the *ssaG* promoter was expressed at a 62% higher level in D33712 than in D23580 confirming the results of the transcriptomic analysis.

Because only a proportion of *S.* Typhimurium cells express certain pathogenicity island-encoded genes during *in vitro* growth (Ackermann *et al*., 2008; Hautefort *et al*., 2003), we determined whether the increased level of expression of SPI2 genes (Fig. 4B) was caused by a higher proportion of D33712 cells expressing SPI2 than D23580 cells. Using derivatives of the two strains that carried the *ssaG*-GFP^+^ construct, we determined the numbers of fluorescent and non-fluorescent cells with flow cytometry (Methods). Under non-inducing conditions, slightly more D37712 cells expressed the *ssaG* SPI2 promoter than D23580 cells (65% vs 60%, respectively) (Fig. 5C). However, this small difference did not account for the 62% increased level of non-induced SPI2 expression seen in Fig. 5B.

SPI2 expression is controlled by a complex regulatory system that operates at both a negative and positive level, involving silencing via H-NS (Lucchini *et al*., 2006), activation by SlyA and SsrB (Fass and Groisman, 2009; Walthers *et al*., 2011) as well as input from OmpR and Fis under non-inducing conditions (Osborne and Coombes, 2011). The reason for the aberrant SPI2 expression in strain D37712 is worthy of further study. Possible explanations include the incomplete silencing of SPI2 transcription or the partial activation of the SPI2 virulence genes under non-inducing growth conditions.

### Increased fitness of *S.* Typhimurium ST313 sublineage L2.2 compared with L2.0 in minimal media

It has become increasingly clear that distinct *Salmonella* pathovariants have evolved particular phenotypic properties that confer fitness advantages during infection of particular avian or mammalian hosts (Branchu *et al*., 2018). Because *S.* Typhimurium ST313 L2.2 appeared to have displaced *S.* Typhimurium ST313 L2.0 in Malawi, we speculated that *S.* Typhimurium ST313 L2.2 might have the competitive edge in some situations. Accordingly, we determined bacterial fitness using a mixed-growth competition assay (Wiser and Lenski, 2015; Lian *et al*., 2023). The competitive index was calculated in three different growth media using pair-wise combinations of strains D37712 and D23580. Two independent approaches were used to phenotypically distinguish the two strains, one based on antibiotic resistance (Fig. 5D) and the other based on fluorescent tagging (Fig. S5).

To confirm that strains engineered to be kanamycin-resistant or gentamicin-resistant did not impact on fitness (Methods), we first verified that the tagged variants of D37712 or D23580 did not confer a growth advantage in LB or NonSPI2 media (Fig. S7). Next, we used a mixed-growth assay to investigate fitness of *S.* Typhimurium ST313 L2.0 strain D23580 or *S.* Typhimurium ST313 L2.2 strain D37712 during growth in LB, or InSPI2 or NonSPI2 minimal media. The data show that both strains grew at similar levels following overnight mixed-growth in nutrient-rich LB media, but D37712 had a competitive advantage during mixed-growth in InSPI2 media (CI = 1.79; *P*<0.05) and a greater competitive edge in NonSPI2 media (CI = 2.20; *P*<0.0001).

We then used an independent fluorescence-based approach to assess the fitness of strains D23580 and D37712 during mixed-growth in NonSPI2 media. This time, the strains were engineered to carry either mScarlet or sGFP2 proteins and the mixed-growth experiments involved pair-wise comparisons of reciprocally-tagged strains. The flow cytometric data showed that in both cases D37712 had a significant competitive advantage in NonSPI2 media (Fig. S5 and S6).

This combination of antibiotic resistance-based and fluorescence-based competitive index experiments lead us to conclude that *S.* Typhimurium ST313 L2.2 strain D37712 had a clear fitness advantage over *S.* Typhimurium ST313 L2.0 strain D23580 during mixed-growth in two formulations of minimal media. The molecular basis of this fitness advantage remains to be established.

### Perspective

Here, we report that *S.* Typhimurium ST313 L2.0 has been clonally replaced by the ST313 sublineages L2.2 and L2.3 as a cause of bloodstream infection in Blantyre, Malawi. In 2018, L2.2 represented the majority of the ST313 strains isolated from hospitalised patients in Malawi at the Queen Elizabeth Central Hospital. Our comparative genomic analysis of ST313 L2.3 identified 30 chromosomal alterations, one of which generated a deletion of the *sseI* effector gene.

Our RNA-seq-based analysis of ST313 L2.2 involved a detailed comparison versus ST313 L2.0 which revealed a key difference involving SPI2 expression. Following initially observations at the transcriptomic level in the ST313 L2 and L2.2 strains grown in a pH-neutral minimal medium (NonSPI2), the increased expression of SPI2 was confirmed at the single cell level using an *ssaG* transcriptional fusion.

A series of experiments showed that the ST313 L2.2 strain D37712 had a competitive advantage over L2 strain D23580 during mixed-growth in minimal media. We propose that this increased fitness of *S.* Typhimurium ST313 L2.2 has contributed to the replacement of ST313 L2.0 in Malawi in recent years.

Previously, we compared three virulence properties of the *S*. Typhimurium ST313 L2.0 D23580 and ST313 L2.2 D37712 strains. First, experiments involving Mucosal Invariant T (MAIT) cells showed that both D37712 and D23580 fail to elicit the high level of activation of MAIT cells that characterises infection by *S*. Typhimurium ST19 4/74 (Preciado-Llanes *et al*., 2020). Second, the D37712 and D23580 strains stimulate similar levels of up-regulation of IL10 gene expression upon infection of human dendritic cells (Aulicino *et al*., 2022). Third, we showed that both D37712 and D23580 express similarly high levels of the PgtE virulence factor that is responsible for the ability of *S*. Typhimurium ST313 to survive human serum-killing (Hammarlöf *et al*., 2018). These findings lead us to conclude that the comparative genomic and transcriptomic differences that distinguish *S*. Typhimurium ST313 L2.0 strain D23580 from ST313 L2.2 D37712 (Fig. 4) do not modulate the ability of the pathogens to activate human MAIT cells or dendritic cells, or to influence the PgtE-mediated serum survival phenotype of *S.* Typhimurium ST313.

Ideally, the implications of the competitive advantage of ST313 L2.2 would be determined in the context of pathogenesis. However, we lack an informative infection model for *S*. Typhimurium ST313 (Lacharme-Lora *et al.,* 2019), and it is not yet possible to experimentally determine whether the improved fitness of L2.2 significantly enhances the success of ST313 during infection of humans.

Here we have investigated the intricate interplay of gene function that is underpinning the success of *S*. Typhimurium ST313 L2.2. We hope that our findings might contribute to future therapeutic or prophylactic strategies for combatting iNTS infections in the African setting.

## Materials and methods

### Bacterial strains

The two *S.* Typhimurium ST313 strains that are the focus of this study are D23580 and D37712. D23580 was isolated from a Malawian 26-month-old child with malaria and anaemia in 2004. D37712 was isolated from the blood of an HIV-positive Malawian male child in 2006. These two African Salmonella strains have been deposited in the National Collection of Type Cultures (NCTC). The D23580 (lineage 2.0) strain is available as <u>NCTC 14677</u>. The ST313 sublineage 2.2 strain D37712 is available as <u>NCTC 14678</u>. All bacterial strains are detailed in Table S8.

### Genome sequencing

The assembled genome and annotation of D23580 (Kingsley *et al*., 2009; Canals *et al*., 2019b) (L2.0) was obtained from the European Nucleotide Archive (ENA) repository (EMBL-EBI) under accession PRJEB28511 (https://www.ebi.ac.uk/ena/data/view/PRJEB28511). For genome sequencing of D37712 (L2.2), DNA was extracted using the Bioline mini kit, and quality was assessed using gel electrophoresis (0.5% agarose gel, at 30 volts for 18 h). The genome was generated by a combination of long read sequencing with a PacBio RS II and short-read sequening on an Illumina HiSeq machine at the Center for Genome Research, University of Liverpool, United Kingdom.

Sequence reads were quality checked using FastQC version 0.11.9 (Andrews, 2010) and MultiQC version 1.8 (Ewels *et al*., 2016), trimmed using Trimmomatic (Bolger *et al*., 2014). Hybrid assembly of the Illumina and PacBio sequence reads was done with Unicycler v0.4.7 (Wick *et al*., 2017).

The assembled genome of S. Typhimurium SDT313 L2.2 strain D37712 was deposited in Genbank (GCA_014250335.1, assembly ASM1425033v1). Raw sequencing reads were deposited for both PacBio and Illumina, under BioProject ID PRJNA656698. Sequence Read Archive (SRA) database IDs are: SRR12444880 for Illumina and SRR12444881 for PacBio.

### Comparative genomic analyses

To generate the data summarised in Fig 1C, sequencing data of 29 *S*. Typhimurium ST313 strains (Msefula *et al*., 2012) were downloaded from EMBL-EBI database (https://www.ebi.ac.uk, accession number ERA015722). Sequence reads were assembled using Unicycler v0.4.8 (Wick *et al*., 2017). The quality of the assemblies was assessed by Quast v5.0.2 (Gurevich *et al*., 2013). The N50 value of all assemblies was >20kb, and the number of contigs was <600.

To construct the phylogenetic tree (Fig 1C), *Salmonella* Typhimurium strains D23580, D37712, LT2 (GCA_000006945.2), DT104 (GCA_000493675.1), 4/74 (GCA_000188735.1), and A130 (GCA_902500285.1) were added as contextual genomes. Roary was used to make the core gene alignment, construct the gene presence/absence matrix and identify orthologous genes (Page *et al*., 2015). Phylogenetic trees were constructed using Randomized Accelerated Maximum Likelihood (RAxML) (Stamatakis *et al*., 2005), and were visualised with the interactive Tree of Life online tool (iToL) (Letunic and Bork, 2006).

The assembled genome and annotation of *S*. Typhimurium ST19 representative strain 4/74 (Richardson *et al*., 2011) were obtained from GenBank (Accession number GCF_000188735.1), while the raw sequencing data of 27 *S*. Typhimurium ST313 strains described in a previous study (Msefula *et al*., 2012) were downloaded from EMBL-EBI database (https://www.ebi.ac.uk, accession number ERA015722). The raw reads were assembled using Unicycler v0.4.8 (Wick *et al*., 2017). The quality of the assemblies was assessed by Quast v5.0.2 (Gurevich *et al*., 2013). The N50 value of all assemblies was >20kb, and the number of contigs was <600.

To identify SNPs, Snippy v4.4.0 (https://github.com/tseemann/snippy) was used to map the raw reads against the 4/74 genome. To detect pseudogene-associated SNPs/indels in each sub-lineage, the SNPs/indels that caused nonsense or frameshifted mutations were filtered. The identifications and names of the disrupted genes were summarised, then the wild type gene sequences were extracted from the 4/74 genome. To validate the pseudogene-associated SNPs/indels, the wild type gene sequences were used to make a BLAST database with BLAST 2.9.0+ (Camacho *et al*., 2009). The 29 genome assemblies were queried against the databases, using the BLASTn algorithm to confirm the nonsense and frameshifted mutations in all isolates.

### Phylogenetic analysis of African *Salmonella* Typhimurium isolates dating from 1966 - 2018

To examine the overall population structure of *Salmonella* Typhimurium responsible for blood infection in Malawi (Fig 1AB and Fig S1), the raw reads of 707 published genome sequences were downloaded (Table S7). Sequence reads were aligned to the *S*. Typhimurium D23580 genome using Snippy v4.4.0. The recombination sites of the alignment were removed by Gubbins (Croucher *et al.,* 2015), and the phylogenetic tree was built with Raxml-ng (Kozlov et al., 2019). The tree was rooted on *Salmonella* Typhi strain CT18 (GCA_000195995.1) as the outgroup. The tree was visualised with the interactive Tree of Life online tool (iToL) (Letunic and Bork, 2006). The sub-lineages were identified with rHierBAPS (Tonkin-Hill *et al.,* 2018). The stacked-area chart and the bar chart showing the percentage and number of isolates from each sub-lineage were made in MS Excel.

### RNA purification and growth conditions

Initially, a screen of transcriptomic gene expression was performed without biological replicates. Total RNA was purified using TRIzol from *S*. Typhimurium D37712 grown in 15 different conditions as described previously (Kröger *et al*., 2013). To generate statistically-robust gene expression profiles, total RNA was subsequently purified using TRIzol from *S*. Typhimurium D37712 grown in four *in vitro* growth conditions (ESP, anaerobic growth, NonSPI2, InSPI2) with three biological replicates as described previously (Kröger *et al*., 2013). RNA was isolated from intra-macrophage D37712 following infection of RAW264.7 murine macrophages using our published protocol (Srikumar *et al*., 2015).

### RNA-seq of *S*. Typhimurium strain D37712 using Illumina technology

For transcriptomic analyses, cDNA samples were prepared from *S*. Typhimurium RNA by Vertis Biotechnologie AG (Freising, Germany). RNA was first treated with DNase and purified using the Agencourt RNAClean XP kit (Beckman Coulter Genomics). RNA samples were sheared using ultrasound, treated with antarctic phosphatase and re-phosphorylated with T4 polynucleotide kinase. RNA fragments were poly(A)-tailed using poly(A) polymerase and an RNA adapter was ligated to the 5’-phosphate of the RNA. First-strand cDNA synthesis was performed using an oligo(dT)-adapter primer and M-MLV reverse transcriptase. The resulting cDNA was PCR-amplified to about 10-20 ng/μl. The cDNA was purified using the Agencourt AMPure XP kit. The cDNA samples were pooled using equimolar amounts and size fractionated in the size range of 200-500 bp using preparative agarose gels. The cDNA pool was sequenced on an Illumina NextSeq 500 system using 75 bp read length.

For the biological replicates of the four growth conditions (ESP, anaerobic growth (abbreviated as NoO2), NonSPI2, and InSPI2) and the intra-macrophage RNA, cDNA samples were generated as above with some improvements in library preparation. First, after fragmentation with ultrasound, an oligonucleotide adapter was ligated to the 3’ end of the RNA molecules. Second, first-strand cDNA synthesis was performed using M-MLV reverse transcriptase and the 3’ adapter as primer, and, after purification, the 5’ Illumina TruSeq sequencing adapter was ligated to the 3’ end of the antisense cDNA. Sequencing of the cDNA was performed as described above. All raw sequencing reads were deposited to the Gene Expression Omnibus (GEO) database under accession GSE161403.

### RNA-seq and dRNA-seq read processing and visualization

RNA-seq data from *S*. Typhimurium 4/74 and D23580 were extracted from previously published experiments (Kröger *et al*., 2013; Srikumar *et al*., 2015; Canals *et al*., 2019b; GEO dataset GSE119724). A combined reference genome was generated that contained the D23580 chromosome plus plasmids pBT1, pBT2, pBT3, pSLT-BT (from D23580) and the D37712 plasmid pCol1B9^D37712^. All reads were aligned and quantified using Bacpipe v0.8a (https://github.com/apredeus/multi-bacpipe). Briefly, basic read quality control was performed with FastQC v0.11.8. RNA-seq reads were aligned to the genome sequence using STAR v2.6.0c using “--alignIntronMin 20 --alignIntronMax 19 --outFilterMultimapNmax 20” options. A combined GFF file was generated by Bacpipe, where all features of interest were listed as a “gene”, with each gene identified by a D37712 locus tag. Subsequently, read counting was done by featureCounts v1.6.4, using options “-O -M --fraction -t gene -g ID -s 1”. For visualization, scaled gedGraph files were generated using bedtools genomecov with a scaling coefficient of 10^9^/(number of aligned bases), separately for sense and antisense DNA strands. Bedgraph files were converted to bigWig using bedGraphToBigWig utility (http://hgdownload.soe.ucsc.edu/admin/exe/linux.x86_64/). Coverage tracks, annotation, and genome sequence were visualized using JBrowse v1.16.6. Transcripts Per Million (TPM) were calculated for all samples and used as absolute expression values (Table S5). A conservative cut-off was used to distinguish between expressed (TPM >10) and not expressed (TPM ≤10), as we previously described (Kröger *et al*., 2013). Relative expression values were calculated by dividing the TPM value for one condition in one strain by the TPM value for the same condition in a different strain. Before the calculation, all TPM values below 10 were set up to 10. A conservative fold-change cut-off of 3 was used to highlight differences in expression between strains.

### Differential gene expression analysis with multiple biological replicates

For differential expression analysis of *S*. Typhimurium strains 4/74, D23580, and D37712, the raw counts (Table S4) from 3-5 biological replicates in four growth conditions were used (ESP, anaerobic growth (abbreviated as NoO2), NonSPI2, and InSPI2). Differential expression analysis was done using DESeq2 v1.24.0 with default settings. A gene was considered to be differentially expressed if the absolute value of its log2 fold change was at least 1 (i.e. fold change > 2), and adjusted p-value was< 0.001.

### The SalComD37712 community data resource, and the associated Jbrowse genome browser

SalCom provides a user-friendly Web interface that allows the visualisation and compaison of gene expression values across multiple conditions and between strains. Particular genes can be selected through pre-defined lists of interest, such as all sRNAs or all genes belonging to a specific pathogenicity island. The resulting heatmap-style display highlights expression differences, and provides access to the rich, manually curated annotation of strains D37712 and D23580. The actual values behind the display can be downloaded for further processing, and a link connects the current view to a genome browser interface.

Visualisation of all the RNA-seq and dRNA-seq (TSS) coverage tracks in JBrowse 1.16.6 shows sequence reads mapped against the combined reference genome described above. Overall, the genomic distance between strains 4/74 and D23580 (approximately 1000 SNPs, or ∼1 SNP per 5000 nucleotides), and between D37712 and D23580 (approximately 30 SNPs, ∼1 SNP per 150,000 nucleotides) allowed the alignment of RNA-seq reads to the simplified combined reference genome without significant loss of reads. The combined reference genome facilitated a direct comparison of gene coverage as well as transcriptional start sites. The unified browser is hosted at http://hintonlab.com/jbrowse/index.html?data=Combo_D37/data.

### Phenotypic and mixed competitive growth experiments

The swimming motility of S. Typhimurium strains D37712, D23580 and 4/74 was determined by a plate assay (Canals *et al*., 2019b), which involved spotting 3 μL overnight culture onto 0.3% LB agar. Relative motility of the three strains was assessed by migration diameter after 4h and 8h of incubation at 37°C.

Relative expression of the *ssaG* SPI2 promoter in strains D23580 and D37712 was measured at the single cell level via GFP fluorescence. Following the construction of a kanamycin-sensitive derivative of D23580 (strain JH4235), a P*ssaG::gfp*^+^ transcriptional fusion was incorporated into the chromosome of JH4235 and D37712 by inserting the *gfp^+^*gene downstream of the *ssaG* gene, under the control of the P*ssaG* promoter. The P*ssaG::gfp*^+^ D23580 derivative (JH4692), and the P*ssaG::gfp*^+^ D37712 derivative (JH4693) are listed in Table S8.

The strains JH4692 and JH4693 were genome sequenced to confirm the integrity of the transcriptional fusions, and to verify that unintended nucleotide changes had not arisen. Following growth in 25 mL non-inducing NonSPI2 media in a 250 mL flask at 37°C with shaking at 220 rpm for approximately 8 hours until OD_600_=0.3, fluorescence was determined with a BD FACSAria Flow Cytometer. The relative fluorescence of the two strains JH4692 and JH4693, and the numbers of individual fluorescent bacteria that expressed the P*ssaG::gfp*^+^ promoter, were determined with FlowJo VX software.

The relative fitness of S. Typhimurium strains D37712 and D23580 was assessed in two independent mixed-growth experiments. First, kanamycin-resistant derivatives of each strain were constructed by inserting the *aph* kanamycin resistance gene into the chromosome at the intergenic region between the *STM4196* and *STM4197* genes, a region that we have previously shown to be transcriptionally silent (Canals *et al*., 2019b). The strains were designated D23580::Km^R^ JH3794 and D37712::Km^R^, JH4232. Mixed cultures of wild-type or kanamycin-resistant derivatives of each strain were grown overnight in LB, InSPI2 and NonSPI2 media in a 250 mL flask at 37°C with shaking at 220 rpm. Following plating on LB agar or LB + kanamycin, colonies were counted and the ratio of bacterial strains was determined. To confirm that the insertion of kanamycin resistance at the intergenic region between *STM4196* and *STM4197* did not impact upon fitness, a mixed-growth experiment was done in both LB and NonSPI2 media (Fig. S7).

Second, to independently assess relative fitness, Tn*7*-based plasmids (Schlechter and Remus-Emsermann, 2019) were used to construct chromosomal sGFP2 and mScarlet derivatives of S. Typhimurium strains D23580 (sGFP2 derivative: JH4694; mScarlet derivative: JH4695) and D37712 (sGFP2 derivative: JH4696; mScarlet derivative: JH4697). The gene cassettes were inserted into the *S.* TyphimuriumTn*7* insertion site between the gene *STMMW_38451* and *glmS*. Mixed cultures of pairs of fluorescently-labelled strains were grown in NonSPI2 media at 37°C with shaking at 220 rpm for approximately 8 hours until OD_600_=0.3. Levels of green and red fluorescence were determined with a BD FACSAria Flow Cytometer.

## Supporting information

**Fig S1.**
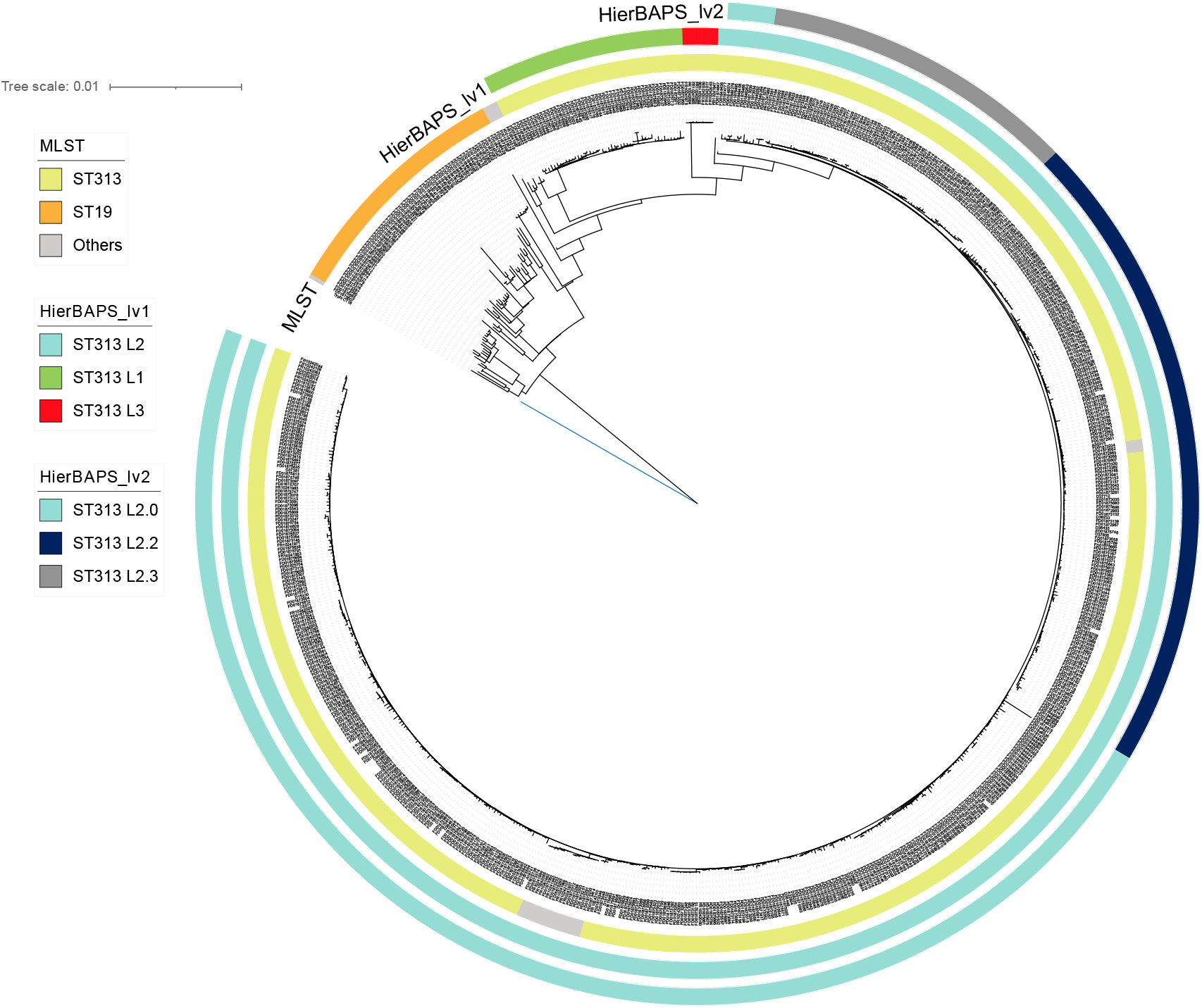
Maximum-likelihood phylogeny of 707 African *S*. Typhimurium isolates. All genome sequences have been published (Msefula *et al.,* 2012, Pulford *et al.,* 2021, Canals *et al.,* 2019b). Raw sequence reads were aligned to the *S*. Typhimurium D23580 genome (FN424405) using Snippy. The recombination sites of the alignment were removed by Gubbins, and the phylogenetic tree was built with Raxml-ng. The tree is rooted on *Salmonella* Typhi strain CT18 as the outgroup. The MLST sequence types, HierBAPS level 1 and level 2 clusters are shown in coloured concentric rings as indicated. The *S*. Typhimurium ST313 isolates are categorised as Lineage 1, Lineage 2 or Lineage 3 according to HierBAPS level 1 clustering. ST313 Lineage 2 was then sub-divided into 3 sub-lineages according to HierBAPS level 2 clustering: ST313 L2.0, ST313 L2.2 and ST313 L2.3. The metadata and lineage designations of all the S. Typhimurium isolates are in Table S7.

**Fig S2.**
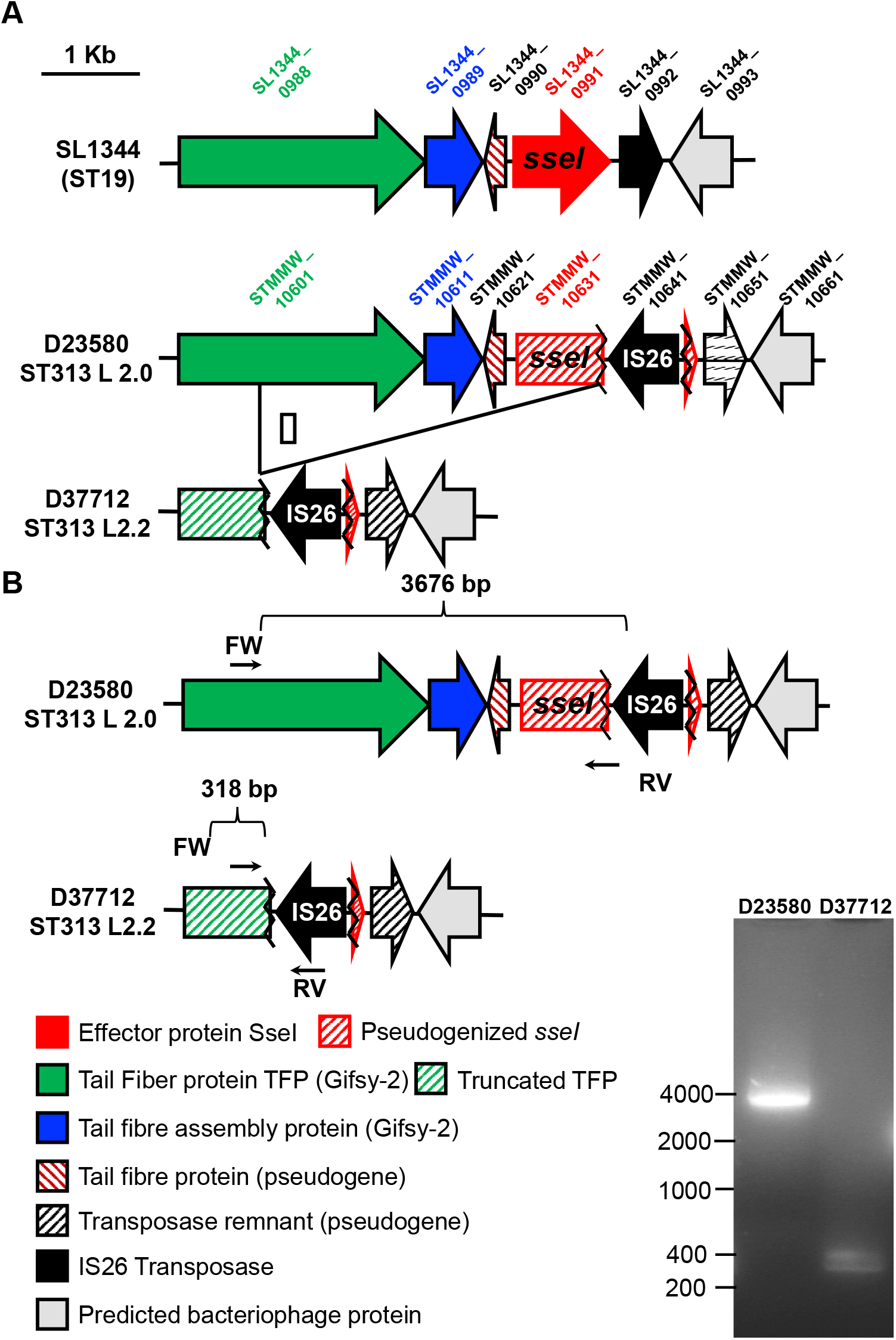
PCR-based confirmation of the deletion of the *sseI* gene from *S.* Typhimurium L2.2 D37712. Arrows from left to right show the forward strand while the left strand is shown by arrows from right to left. However, *sseI* gene in D23580 is a pseudogene with a SNP indicated as a red line.

**Fig S3.**
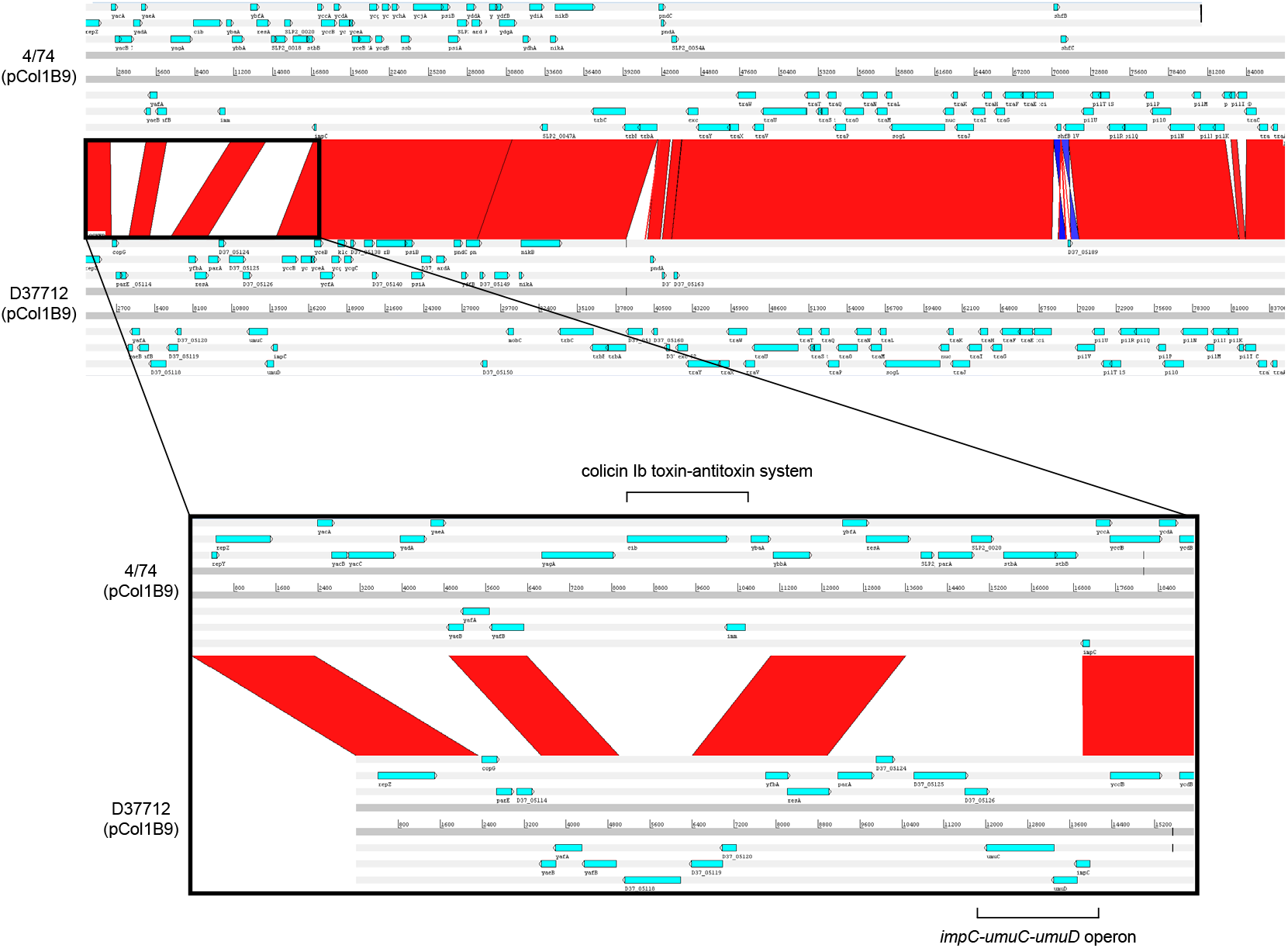
Genomic comparison of plasmids pCol1B9^4/74^ and pCol1B9^D37712^ using Artemis Comparison Tool (ACT). Bottom panel details the differences observed in the most divergent regions, including colicin toxin-antitoxin system (in pCol1B9) and *impC-umuC-umuD* operon (in pCol1B9).

**Fig S4.**
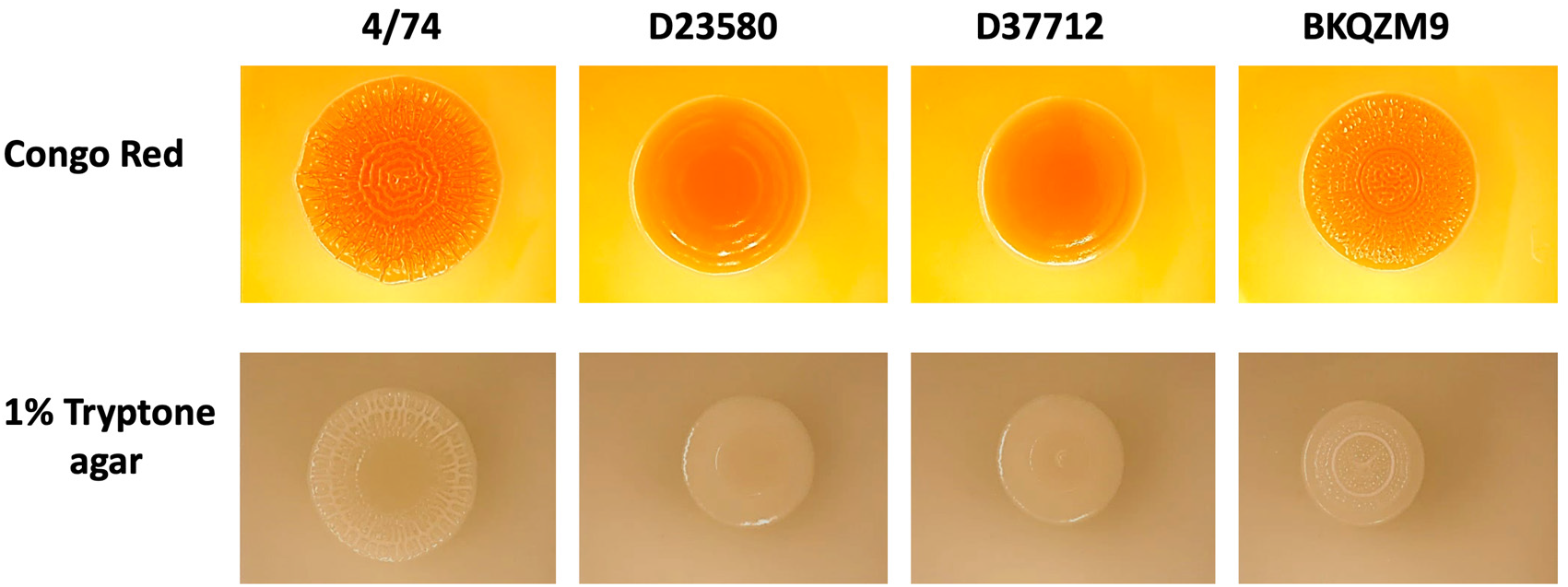
RDAR Phenotypes of 4/74, D23580, D37712 and BKQZM9. The top panel shows the RDAR morphology assay and the bottom panel shows a complementary experiment that involves the induction of biofilm formation on 1% tryptone agar (MacKenzie *et al.,* 2019). Strain 4/74 was used as a RDAR-positive control, which has concentric rings and a wrinkled appearance (Pulford *et al.,* 2021). The S. Typhimurium ST313 L3 strain BKQZM9 is shown for comparative purposes.

**Fig S5.**
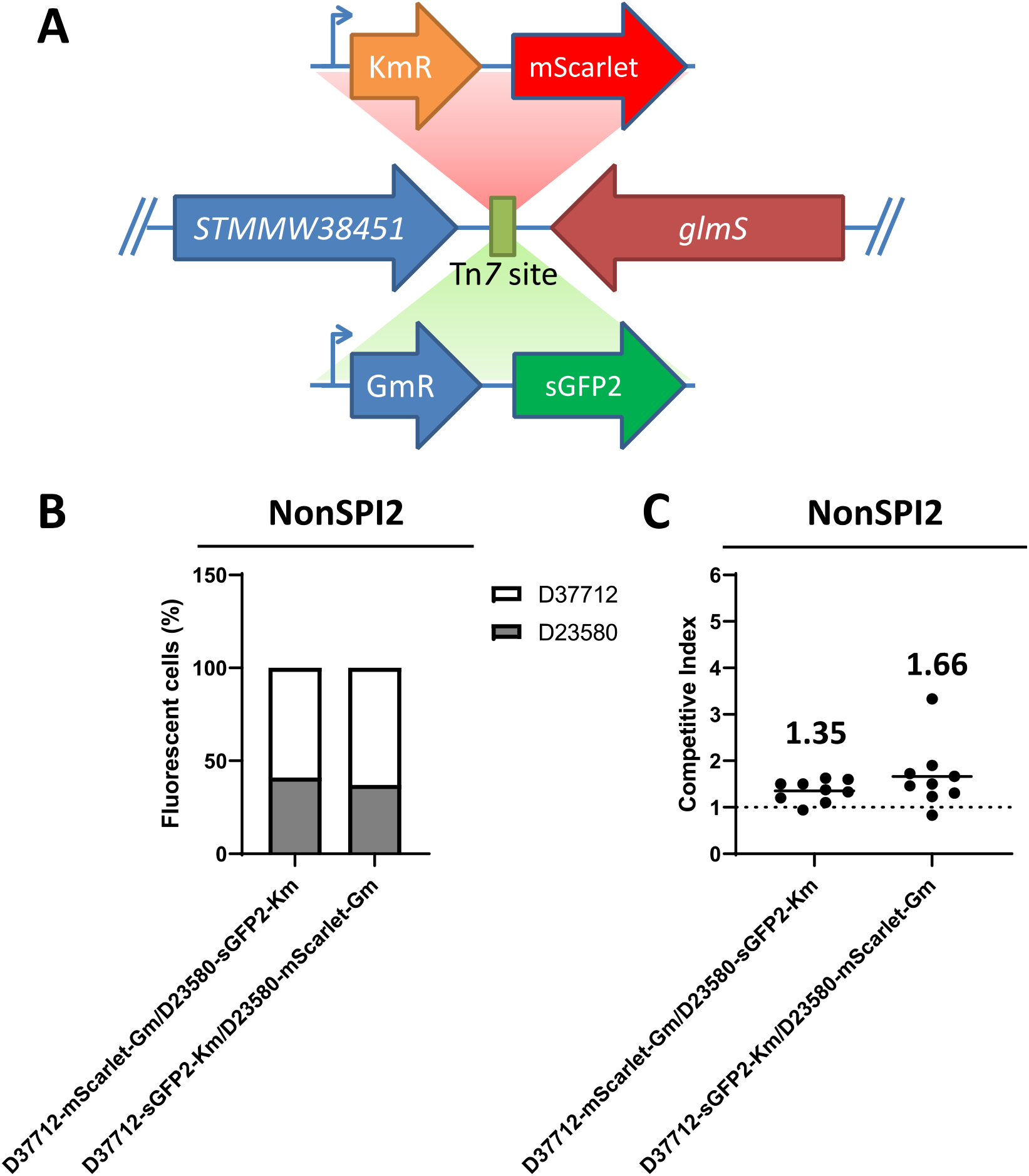
Competitive index analysis of D23580 and D37712 using fluorescently-tagged S. Typhimurium strains. **(A)** Km^R^-sGFP2 and Gm^R^-mScarlet were inserted into the transposon Tn7 site of D23580 or D37712. Bent arrows represent promoters and directional arrows represent genes. **(B)** A 1:1 mix of Km^R^-sGFP2 and Gm^R^-mScarlet marked strain was inoculated in NonSPI2 media, followed by an overnight incubation in 37℃. Percentage of sGFP2 (green) and mScarlet (Red) marked cells was measured by flow cytometry. Raw data are shown in Figure S7, 10,000 events were acquired for each sample. **(C)** Competitive index analysis of Km^R^-sGFP2 and Gm^R^-mScarlet marked strain. Bacterial numbers were determined by counting CFU for overnight culture of a 1:1 mixture in NonSPI2 media. Each dot represents a single biological replicate and the lane represents mean value. A competitive index of 1 indicates the equal fitness of two strains, while a number higher than 1 reflects an increased fitness of D37712.

**Fig S6.**
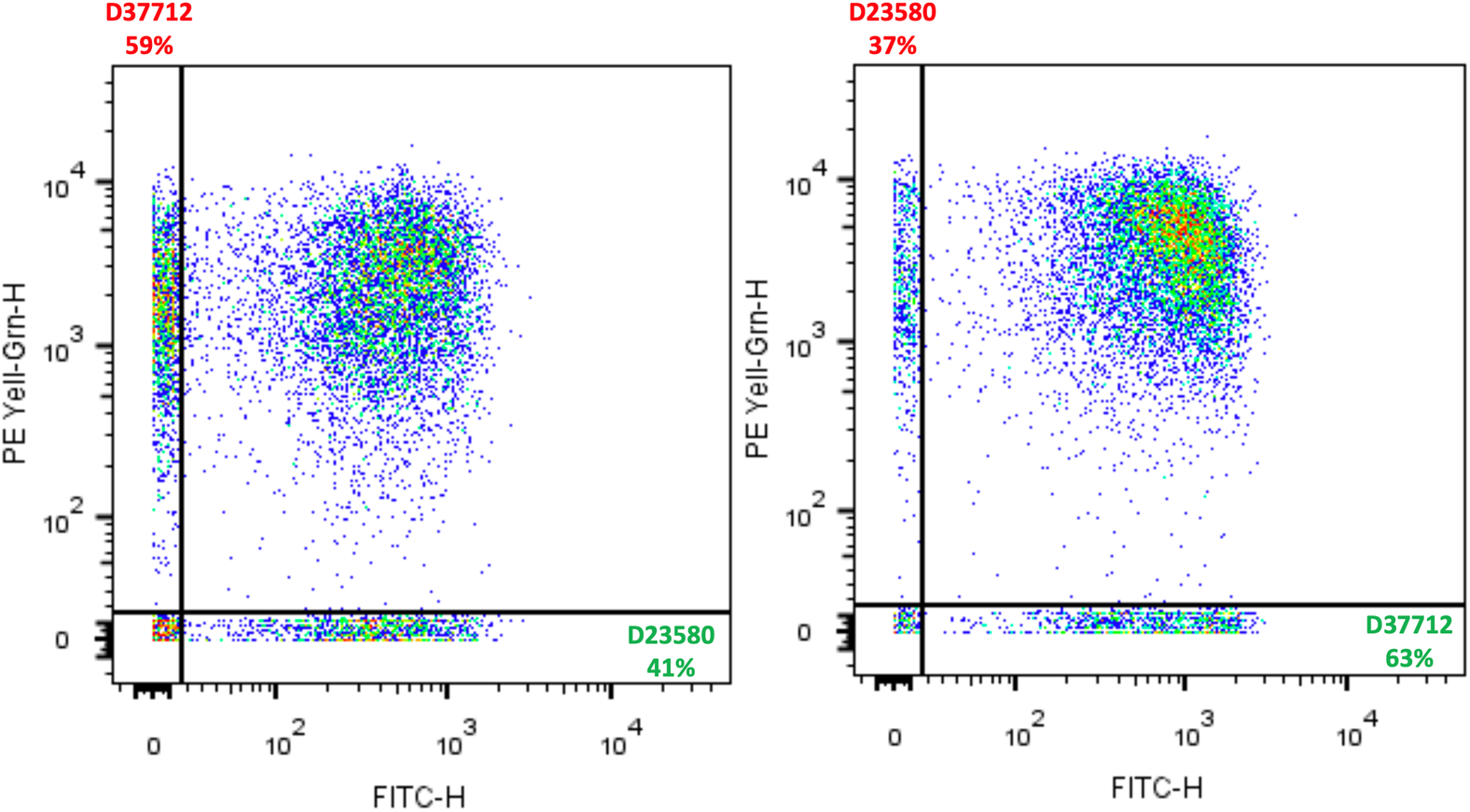
Raw flow cytometric data related to Fig. S5B. **(A)** JH4695 + JH4698 and **(B)** JH4696 + JH4697. A 1:1 mix of the Km^R^-sGFP2 and Gm^R^-mScarlet marked strains were inoculated in NonSPI2 media, followed by growth at 37℃ until OD_600_ = 0.3. The X-axis (labelled FITC) shows the GFP level and the Y-axis (labelled PE Yell-Grn) indicates the mScarlet level. Quadrant gates were used to separate four populations, and the black numbers indicate the percentage of events in each quadrant. In total, 10,000 events were acquired for each sample.

**Fig S7.**
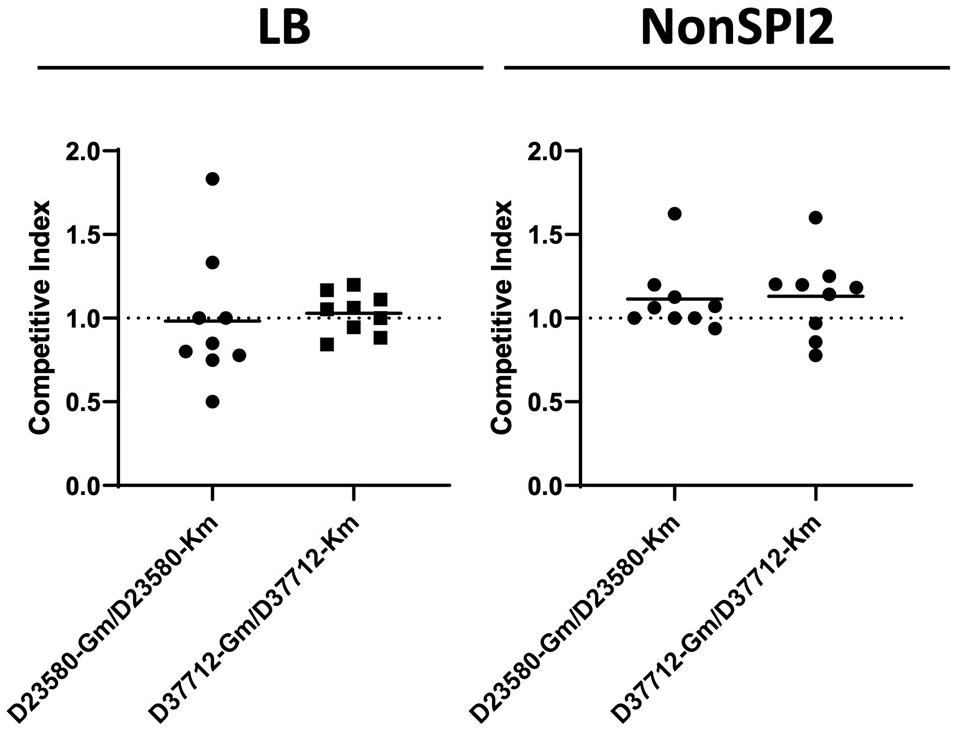
The insertion of GFP-Km or RFP-Gm did not impact on fitness. A 1:1 mix of Km^R^-sGFP2 and Gm^R^-mScarlet marked strains were inoculated in LB or NonSPI2 media, followed by overnight incubation in 37℃. The competitive index (CI) was calculated using the formula (CFU_Gm)_/(CFU_Km_). Each dot represents the CI from a single replicate and the horizontal bars indicate the mean of each dataset.

## Supplementary data

**Table S1:** SNP and indel variants that differentiate L2.2 (strain D37712) and L2.3 (strain D49679).

**Table S2:** SNP and indel variants that differentiate L2.2 (strain D37712) and L2.0 (strain D23580).

**Table S3:** Pseudogenes carried by ST19 and ST313 L2.0 and L2.2 (strains 4/74, D23580 and D37712).

**Table S4:** Raw read counts for all processed RNA-seq samples shown in Figures 3 and 4 (strains 4/74, D23580, and D37712).

**Table S5:** TPM values for all processed RNA-seq samples shown in Figures 3 and 4 (strains 4/74, D23580, and D37712).

**Table S6:** Differential expression analysis using DESeq2 for strains D23580 vs D37712 grown in four *in vitro* conditions.

**Table S7:** Metadata and lineage designations of the 708 S. Typhimurium isolates used to generate the maximum likelihood phylogeny (Fig. S1).

**Table S8:** Bacterial strains used in this study.

## Acknowledgements

We are grateful to present and former members of the Hinton laboratory for helpful discussions, and to Paul Loughnane for his expert technical assistance.

This work was supported by a Wellcome Trust Investigator award [grant numbers 106914/Z/15/Z and 222528/Z/21/Z] to J.C.D.H., and by the Malawi-Liverpool-Wellcome Research Centre Director’s Fund. B.K. was funded by an AESA-RISE fellowship from the African Academy of Sciences [Grant Number: RPDF-18-04]. For the purpose of open access, the authors have applied a CC BY public copyright licence to any Author Accepted Manuscript version arising from this submission.

## Author contributions

**Conceptualization:** B.K., RH, M.A.G., C.L.M. and J.C.D.H.

**Data curation:** B.K., R.C., A.V.P., C.V.P., P.A.

**Formal analysis:** B.K., R.C., C.V.P., A.V.P., X.Z., C.K., S.V.O., Y.L., P.A., A.D. and J.C.D.

**Funding acquisition:** B.K., R.C., A.V.P., X.Z. and J.C.D.H.

**Investigation:** B.K., R.C.A., A.V.P., X.Z. and J.C.D.H.

**Methodology:** B.K., R.H., M.A.G., C.L.M. and J.C.D.H.

**Project administration:** B.K. and J.C.D.H.

**Resources:** B.K., R.H., M.A.G, C.L.M. and J.C.D.H.

**Software:** B.K. and A.V.P.

**Supervision:** M.A.G., C.L.G., C.L.M. and J.C.D.H.

**Validation:** B.K., R.C., A.V.P., X.Z. and J.C.D.H.

**Visualization:** B.K., R.C., Y.L., C.V.P., A.V.P. and J.C.D.H.

**Writing original draft:** B.K., R.C. and J.C.D.H

**Writing reviews and editing:** B.K., R.C., A.V.P., X.Z., C.K., S.V.O., A.D., R.H., M.G and J.C.D.H.

**Equal contribution:** Authors B.K., R.C. and A.V.P. made equal contributions to this work.

## Notes

http://hintonlab.com/jbrowse/index.html?data=Combo_D37/data

### Competing Interest Statement

The authors have declared no competing interest.

